# Brain2Pix: Fully convolutional naturalistic video reconstruction from brain activity

**DOI:** 10.1101/2021.02.02.429430

**Authors:** Lynn Le, Luca Ambrogioni, Katja Seeliger, Yağmur Güçlütürk, Marcel van Gerven, Umut Güçlü

## Abstract

Reconstructing complex and dynamic visual perception from brain activity remains a major challenge in machine learning applications to neuroscience. Here we present a new method for reconstructing naturalistic images and videos from very large single-participant functional magnetic resonance imaging data that leverages the recent success of image-to-image transformation networks. This is achieved by exploiting spatial information obtained from retinotopic mappings across the visual system. More specifically, we first determine what position each voxel in a particular region of interest would represent in the visual field based on its corresponding receptive field location. Then, the 2D image representation of the brain activity on the visual field is passed to a fully convolutional image-to-image network trained to recover the original stimuli using VGG feature loss with an adversarial regularizer. In our experiments, we show that our method offers a significant improvement over existing video reconstruction techniques.

## Introduction

A long-lasting interest of sensory neuroscience is understanding how sensory information is represented in neural activity patterns. Decoding visual stimuli from neural activity using deep learning is a promising approach for bringing us closer to such understanding. Recent advances allow the successful decoding of static images from brain data [5, 10, 45, 30, 32, 17, 25, 12, 41]. Reconstructing natural movies is significantly more challenging [34] yet important given that neurons respond to signals that unfold over both space and time [33]. The difficulty with reconstructing natural movies is in large part due to the limited temporal information provided by imaging methods such as fMRI as well as the complex dynamics of the natural world that the model must learn.

Convolutional image-to-image models have recently achieved unprecedented results in multiple tasks such as semantic segmentation [29, 36, 35, 28, 50], style transfer [52, 11, 42, 22], colorization [48, 20, 49] and super-resolution [24, 6, 51]. Convolutional image-to-image networks have the great advantage of preserving the topography of input images throughout all the layers of the network. Consequently, the network does not need to learn a remapping between locations and can focus on processing local features. The reconstruction of perceived natural images from brain responses can be considered as a form of image-to-image problem since the visual cortex processes information in a topographically organized manner [14, 15, 21] such that the topology of the input images is preserved within each visual area. The retinotopic mapping of visual neurons defines relationships between the visual field and its cortical representation in individual subjects and has uncovered many important aspects of the visual cortex across different species [18, 7]. However, it is not straightforward to exploit this in an image-to-image ConvNet architecture. The cortex itself can be roughly seen as a pair of topological spheres embedded in a 3D space. Several separate visual representations are embedded in this cortical space, corresponding to several visual areas (e.g. V1, V2, V3). These representations are furthermore distorted by the geometry of the cortex and by the uneven sampling of different parts of the visual field. Therefore, there is no natural way of constructing a convolutional architecture that exploits the image-to-image nature of the problem by preserving the topography between voxel responses and pixel brightness and color.

In this paper, we exploit the receptive field mapping of visual areas to convert voxel responses defined in the brain to activations in pixel-space. Early visual areas V1, V2 and V3 were identified using retinotopy. The voxel activations of each area are then converted to images via the mapping of the receptive fields. Importantly, these images (visual representations) do have a pixel-to-pixel correspondence with the images used as stimuli. We then transform these visual representations into realistic images using an image-to-image U-network trained using a combination of pixelwise, feature and adversarial losses.

## Related work

Recent work on image reconstruction from fMRI data has demonstrated the success of employing deep neural networks (DNNs) and generative adversarial networks (GANs) in neural decoding [34, 16, 12, 47, 41, 13, 43, 44]. For instance, [41] used a GAN to reconstruct grayscale natural images as well as simpler handwritten characters. Another approach was voxel-wise modeling by [34], where they modeled responses to complex natural stimuli to estimate the semantic selectivity of voxels. More recently, [43] showed that even with a limited set of data – in the order of thousands compared to millions that the field is accustomed to – it was possible to train an end-to-end model for natural image stimulus reconstruction by training a GAN with an additional high-level feature loss. Their reconstructions matched several high-level and low-level features of the presented stimuli. However, a comparable performance has not yet been achieved for naturalistic video stimuli. The most recent notable video reconstruction study by [13] made use of a variational autoencoder and was able to reconstruct low-level properties of the images, where the reconstructions resembled shadows or silhouettes of the stimulus images. Reconstruction of perceived videos can thus be considered a very challenging problem.

The simplest way to apply convolutional neural networks (ConvNets) on fMRI voxel responses is to treat fMRI slices as separate images stacked on the channel dimension [38]. However, these images do not respect the topography of neural representations and contain a large fraction of non-responsive voxels corresponding to white matter and cerebrospinal fluid. This results in most of the contrast of the images depending on irrelevant anatomical factors. Another possibility is to use spatial 3D convolutions on the brain volume [1]. This method has the benefit of preserving the topography of the neural responses but otherwise has the same issues as the 2D approach. These shortcomings make such methods unsuitable for brain decoding and reconstruction. A more viable strategy is to map the voxel responses on a mesh representing the cortical surface [9] and apply a geometric deep learning technique [31, 8, 4, 26].

## Material and Methods

Our brain2pix architecture has two components: 1) a receptive field mapping that transforms the brain activity of visual regions to a tensor in pixel (input) space, exploiting the topographical organization of the visual cortex; 2) a pix2pix network that converts the brain responses in pixel space to realistically looking natural images. In the following, we describe the two components in detail.

### From voxels to pixels

A receptive field mapping is a (potentially many-to-one) function that maps the 3D coordinate of the voxels of a visual area to Cartesian coordinates in the stimulus space. This coordinate is defined as the region of the image that elicits the highest response in the voxel. Given a visual ROI, we can refer to these mappings using the following notation:

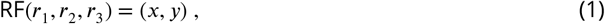

where (*r*_1_, *r*_2_, *r*_3_) are the voxel coordinates and (*x, y*) are Cartesian coordinates in stimulus / pixel space in the image space. Since visual areas are topographically organized, this map can be seen as an approximate homeomorphism (i.e. a function that preserves the topology). Note that RF does not respect the metric structure of the image since the representation of the fovea is inflated while the periphery is contracted. We denote the function associating a measured neural activation (BOLD response) to each voxel as *n*(*r*_1_, *r*_2_, *r*_3_). Using the receptive field mapping, we can transport this activation map to pixel space as follows:

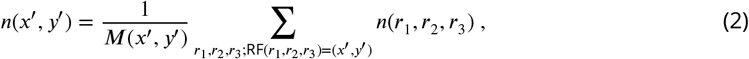

where *M*(*x′*, *y′*) is the number of voxels that map to the coordinates (*x′*, *y′*). Eq. 1 is limited to the case of point-like receptive fields. More generally, the RF transport map can be written as a linear operator:

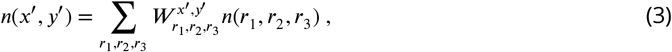

where the weight tensor *W* is a (pseudo-)inverse of the linear response function of the cortex under single pixel simulations. This second formulation has the benefit of allowing each voxel to contribute to multiple pixels and to be suitable to gradient descent training.

In this paper we use two strategies for determining *W*. The first approach, is to apply an off-the-shelf receptive field estimator and to use Eq 1. The second, more machine learning oriented approach, is to learn a weight matrix together with the network. In order to preserve the topographical organization, we include the learnable part as a perturbation of the receptive field estimation:

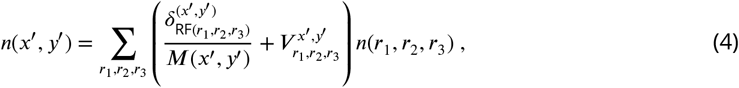

where 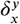 is the discrete delta function and the weights 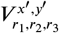 are learnable parameters.

### Image-to-image network

The input to the pix2pix network is a tensor obtained by stacking the voxel activation maps, one map for each combination of ROI and time lag. In fact, the network needs to integrate the topographically organized information contained in several layers of the visual hierarchy (V1, V2 and V3 in our case) but also the responses at different time lags (the network selects from the 5 provided RFSimages).

#### Architecture

The architecture of the brain2pix model is inspired by the pix2pix architecture [22] which comprises a convolutional U-Net-based generator [36] and a convolutional PatchGAN-based discriminator (Figure 1). The first and the last layers of the generator are respectively convolutional and deconvolutional with four standard U-net skip blocks in-between. All five layers of the discriminator are convolutional with batch normalization and a leaky ReLU activation function.

**Figure 1.**
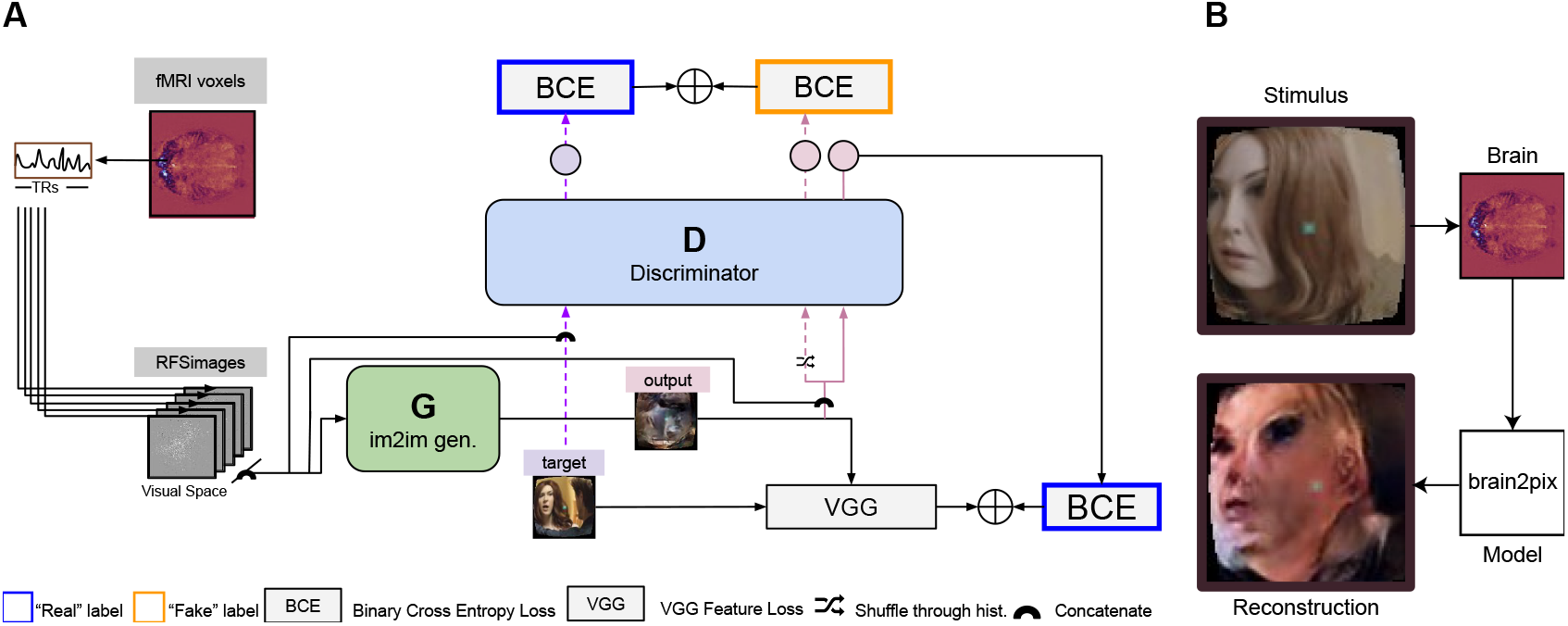
A) Illustration of the brain2pix model. First, the voxels of each region of interest (ROI) voxels are extracted. The voxel receptive fields are then projected onto the input (visual) space. Using this mapping the RSFimages are created, which reflect voxel activity at the locations of their receptive fields in the visual space. To account for the hemodynamic delay, these activities are taken with a fixed delay of 5 TR (i.e. the activities are taken from the volume recorded approximately 4 seconds after stimulus presentation). The input to the **generator** are the RSFimages, and the outputs are the reconstruction (end-to-end modeling). The loss is estimated between the target (original) image and this reconstruction as a feature loss (VGG loss). The reconstruction is also passed through the discriminator, concatenated with the RFSimages. The BCE loss estimated on the discriminator’s output is summed with the VGG loss, and this sum of losses is then backpropagaged to update the parameters of the generator. The loss for training the **discriminator** is obtained by summing its output based on the reconstructed image and the target image (concatenated with the RFSimages), using a BCE loss. B) Example reconstruction on the test set. The end-to-end model reconstructs static video frames, using the brain signal of the participant watching Doctor Who in the fMRI scanner as the input.

The discriminator was trained to distinguish stimuli from their reconstructions by iteratively minimizing a loss function with a sole adversarial loss (binary cross-entropy (BCE)) and using a history buffer to encourage the discriminator to remember past errors.

The generator was trained for converting brain responses to stimulus reconstructions by iteratively minimizing a loss function with three weighted components: i) pixel-loss, which was taken to be the absolute difference between ground-truths and predictions, ii) feature loss, which was taken to be the Euclidean distance between pretrained layer 10 VGG features of ground-truths and predictions and iii) adversarial loss, which was taken to be the “inverse” of the adversarial loss that was used to train the discriminator.

All models were implemented in Python with the MXNet framework [3]. They were trained and tested on Nvidia GeForce 2080 Ti GPUs.

#### Receptive field estimation

Receptive fields for dorsal and ventral visual regions V1, V2 and V3 were estimated in a data-driven way using *neural information flow* [39]. Grayscale video sections were passed through three 3D convolutional neural network layers corresponding to the visual ROIs. Before the ROI-specific layers a linear layer with a single 1 × 3 × 3 channel was used to allow learning retinal and LGN preprocessing steps. Average pooling was applied after each layer to account for increasing receptive field sizes, the temporal dimension was average pooled to a TR of 700ms before applying the observation models, and spatio-temporal receptive fields were constrained to be positive. For training this neural network, a low-rank tensor decomposition was applied to estimate voxel-wise spatial, temporal and channel observation (readout) vectors, which were used to predict voxel-wise activity from the neural network activity tensors. The receptive field location (*x, y*) for every voxel was then estimated as its center of mass of the low-rank receptive field maps.

### Data acquisition

We made use of a public large fMRI dataset from single-participant responses to naturalistic video stimuli [40]. The exact experiments are described in detail in the original study [40]. In short, the participant fixated the center of the screen and watched 30 episodes of BBC’s Doctor Who while their BOLD activity was measured. The videos were presented in multiple sessions, using a head cast for positional stability and consistency. The recording comprised presenting 30 full episodes once (forming the training set, here used for model estimation), and 7 short clips – teasers and short stories, taken from a different Doctor to avoid train-test overlap – repeatedly shown at the end of most sessions (forming the test set). The 22–26 repetitions of the test set were averaged and used for evaluation of the brain2pix model. The averaged test set is common in neural coding and most reconstruction research to provide a version of the individual’s BOLD response patterns with high signal-to-noise ratio, freed of the substantial noise contained in fMRI data. To assess the implications of reconstruction research it is important to understand that the presented reconstructions have so far usually been estimated on data from specific individuals averaged over multiple presentations in highly controlled settings.

### Data preprocessing

Prior to using the inputs for training the model, 3D brain matrices were transformed to 2D receptive field signal images (RFSimages) in two main steps. First, regions of interests (ROIs) were selected from the brain (V1, V2, V3), based on their corresponding masks. Second, voxels in each brain region were mapped onto their corresponding visual space based on the retinotopic map. The 2D RFSImages were resized to 96 × 96 pixels and separated by five time channels (TRs) and amount of brain regions (V1, V2, V3).

The videos were downsampled spatially (96 × 96 × 3) and temporally to match the TRs of the fMRI recordings (one frame every 0.7 s). This resulted in a total of approximately 119.000 video frames for training and 1034 video frames for model evaluation. To incorporate the haemodynamic delay we realigned the stimuli and brain signals such that the current signals correspond to the stimuli that were presented at 3 TRs before, allowing a time window of 2.1 s – 4.2 s delay from stimulus presentation (since we are incorporating 3 TRs). Finally, each frame underwent a fish-eye transformation, which mimics biological retinal eccentricity [2]. The receptive field centers we used for mapping brain signals onto the visual space were based on images that underwent this transformation.

### Experimental design

We compared our final model with alternative reconstruction models. This included a baseline comparison where brain2pix was compared with previously suggested models. Since we wanted to focus on early visual areas, we also trained our model on V1, V2, and V3 individually (which we called the ROI experiment). Finally, we tested whether our model was robust to various ablations.

The same four evaluation metrics were used in all experiments, namely Pearson’s product-moment correlation coefficient (corr.) and Euclidean distance (dist.) between the features of the presented test stimuli and their reconstructions. Features were extracted from the pool2, pool5 and fc6 layers of the AlexNet model [27] and the C3D model [46]. The Alexnet model was pretrained on ImageNet [37] and the c3d model was pretrained on Kinetics-400~[23]. Additional details of the experiments, additional results and a link to the source code are provided in the supplementary materials.

## Results

### Experiments

This paper consists of 7 experiments: 1) The brain2pix architecture trained on synthetic data with fixed receptive field locations, 2) training brain2pix using real fMRI data with a fixed receptive field (Fixed RF) for reconstruction, 3) training brain2pix on real data with learnable receptive fields (LearnedRF) for reconstruction, 4) training traditional models for comparison with the brain2pix model (baseline experiment), 5) training brain2pix on fMRI data from various brain regions with fixed RF (ROI experiment), 6) removing essential components from the brain2pix architecture with fixed RF (ablation experiment) and 7) experimenting with various sizes of training data with a fixed RF.

### Brain2Pix on synthetic data

Before experimenting on real data, we used synthetic data to test feasibility and tune hyperparameters of brain2pix. Instead of mapping BOLD reponses for each ROI and voxel onto the pixel space, we mapped target stimuli onto pixel space using the same exact method. This filtering of target images with the RF centers gave us the same amount of input pixels for the model, all at the same location. However, their activations were not based on actual brain signals, but rather on the pixels of the target images, which resulted in the same number of pixels as the number of voxels provided by the three brain regions used. We got clear reconstruction images from this, which confirmed that the number of RF pixels provided by V1 + V2 + V3 ROIs could theoretically carry enough spatial information for the model to generate realistic and accurate results.

### Brain2Pix variants

The brain2pix variants described in detail below differed in how they transformed brain responses from volumetric representation to image representation.

#### FixedRF

The protocol for our main model is to use the (fixed) receptive field estimates. Once the brain signals were mapped onto visual space and the model was assembled, we ran it to obtain reconstructions (see Figure 2 and Figure 3 under Brain2pix > Fixed RF). The results show individual frames from a snippet of the fMRI test set. The figure contains a selection of frames from the test set and their corresponding reconstructions. This figure shows that the model successfully reconstructed frames that contained head shapes, silhouettes, facial expressions, and also objects (such as a white blanket in frames 289-292). The figure also depicts a smooth transitioning between frames, which allows better reconstruction of video clips. Table 1 and Table 2 contain the quantitative results of the model under “B2P-FixedRF”. This model achieves the highest performance in terms of correlation for most of the studied feature layers. However for the distance values, it shows the best results for only one layer.

**Figure 2.**
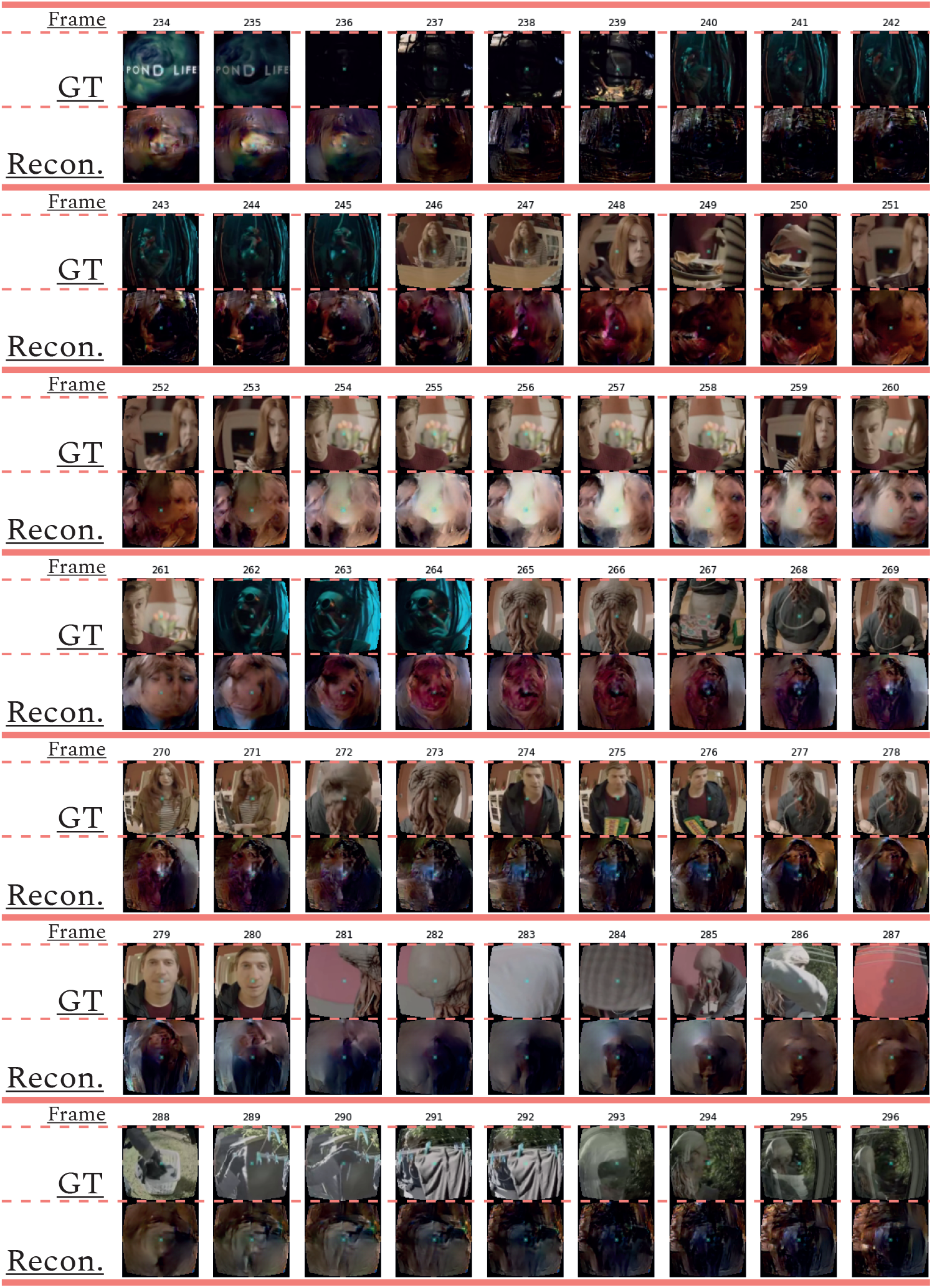
Sequential of reconstructed frames: Consecutive frame sequence from a video fragment of the test data (ground truth (GT)) and the corresponding reconstructions (Recon.).

**Figure 3.**
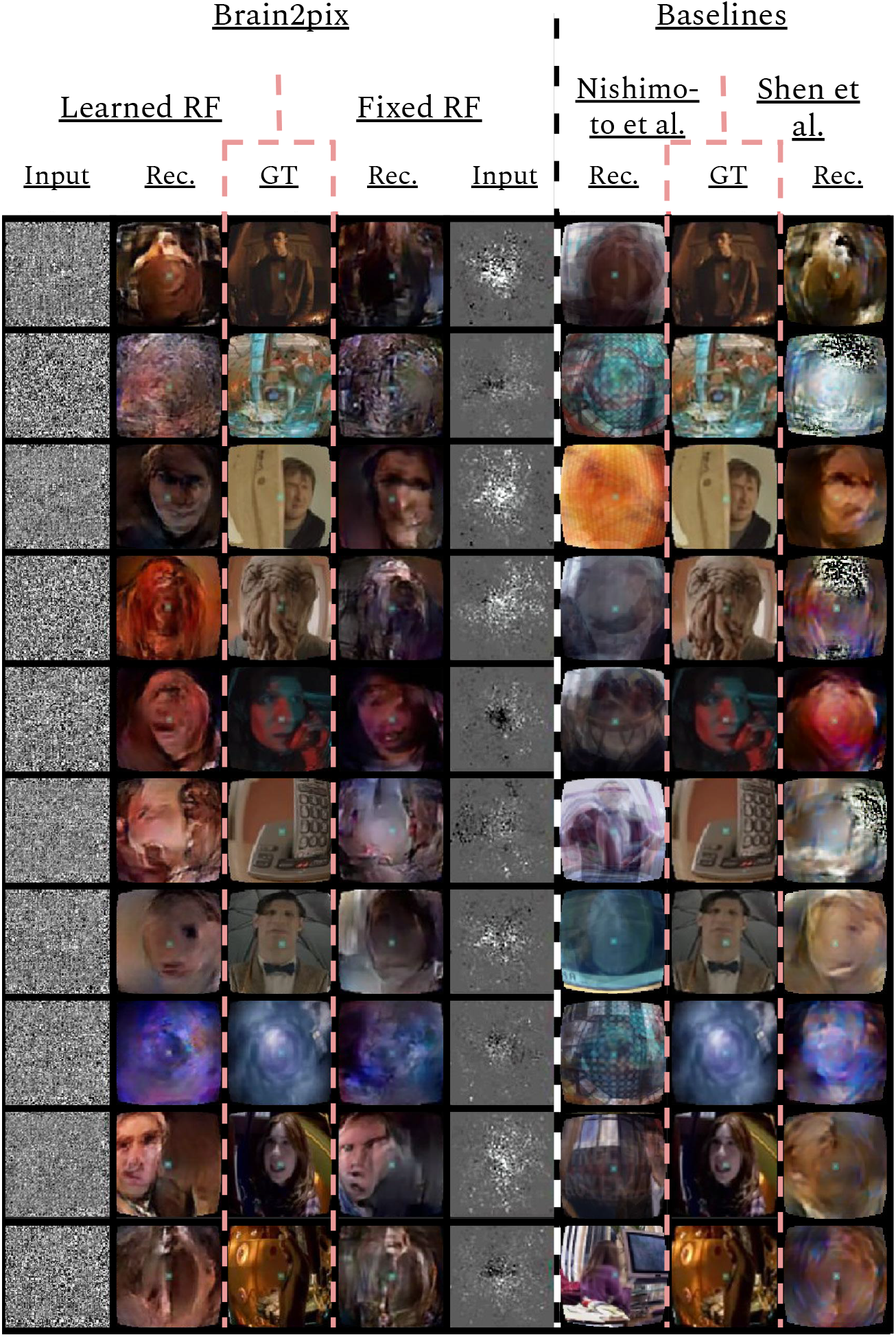
Baseline experiment: Comparison between the reconstructions of the brain2pix and the baseline models. The reconstructions of brain2pix are shown in columns 2 and 4, with their corresponding ground truth (GT) in the middle column 3, and inputs in columns 1 and 5. The reconstruction of the baseline models are shown in columns 6 and 8 together with their corresponding GT in the middle column 7.

**Table 1.** Baseline experiment *correlation values*: Correlation values between the features of the reconstructions of the baseline experiment and features of the target taken from various layers obtained from passing target / reconstructions through a pretrained network (Alexnet and C3D). Here comparisons are done for the brain2pix models and baseline models. Bold values indicate the highest correlation values.

|  | B2P-learnRF | B2P-fixedRF | Nishimoto et al. | Shen et al. | FC-layers |
| --- | --- | --- | --- | --- | --- |
| AlexNet pool2 | 0.4573 | **0.4615** | 0.2526 | 0.4141 | 0.0479 |
| AlexNet pool5 | **1.3983** | 0.3507 | 0.2318 | 0.3269 | -0.0394 |
| AlexNet fc6 | **0.4653** | 0.4608 | 0.2016 | 0.4196 | -0.0214 |
| C3D pool2 | 0.4831 | **0.4869** | 0.3662 | 0.4212 | 0.1284 |
| C3D pool5 | 0.2356 | **0.2426** | 0.0489 | 0.2030 | 0.0427 |
| C3D fc6 | 0.2426 | **0.2519** | 0.0412 | 0.2185 | 0.0434 |

**Table 2.** Baseline experiment *distance values*: Distances between reconstruction and target features of various layers obtained from passing reconstructions / targets through a pretrained network (Alexnet and C3D). Here comparisons are done for the brain2pix models and baseline models. Bold values indicate the lowest distance values.

|  | B2P-learnRF | B2P-fixedRF | Nishimoto et al. | Shen et al. | FC-layers |
| --- | --- | --- | --- | --- | --- |
| AlexNet pool2 | **4.6089** | 4.6252 | 5.5266 | 5.3511 | 56.8820 |
| AlexNet pool5 | **1.2425** | 1.2692 | 1.2432 | 1.2532 | 3.3593 |
| AlexNet fc6 | 1.4062 | **0.0899** | 1.7322 | 1.4861 | 5.5175 |
| C3D pool2 | **1.3842** | 1.3983 | 1.4238 | 1.8132 | 31.6262 |
| C3D pool5 | 3.4850 | 3.4626 | **2.6601** | 3.8881 | 9.6941 |
| C3D fc6 | 0.9269 | 0.9332 | **0.8655** | 1.1847 | 4.8835 |

#### LearnedRF

The second variant (referred to as Learned RF) used a dense layer to perturb the image representation in the Fixed RF variant as a function of the volumetric representation in order to imrpove it even further. Reconstructions of this model are shown in Figure 3 under Brain2pix > LearnedRF. We found that the LearnedRF variant has the highest performance in terms of correlation for the Alexnet pool2 and Alexnet fc6 layers (see Table 1 and Table 2).

The differences in the quantitative and qualitative results between LearnedRF and FixedRF were not large, suggesting that both the RF models capture the correct topographical structures. Both brain2pix variants contain significantly above chance level performance (p < 0.05; Student’s t-test) and significantly outperformed both baselines (p < 0.05; binomial test) (Figure 3).

### Baselines

#### Nishimoto Baseline

We compared our brain2pix results with baseline models based on state-of-the-art reconstruction models. The first baseline is based on the method introduced by Nishimoto et al., which used a set of naturalistic videos as an empirical prior [32, 34]. In our experiment, we used a smaller but more targeted natural image prior constructed from the training set. In short, we trained an encoding model, consisting of a dense layer, that predicts BOLD activity from c3d features extracted from training samples, which we used as the likelihood. Then we constructed the images by averaging the 10 clips with the highest likelihood.

The reconstructions resulting from this method are shown in Figure 3 in the columns under the header “Baselines > Nishimoto et al. > Rec.”. Unlike our method, this method did not make use of retinotopic information, and could be overfitted to the training image distribution. This resulted in difficulty of the model in reconstructing things that are not present in the training set, such as the Ood character. Additionally, some reconstructions do not come close to the target perceptually, such as the 10th row in Figure 3 where a computer is reconstructed although it is supposed to be a person. Quantitative results are shown in Table 1 and Table 2, which indicates all lower correlation values and mainly higher distances (except for C3D pool5 and C3D fc6) compared to our model. The correlations (performance estimation) are generally lower than the Shen baseline, but higher than a model with only FC-layers.

#### Shen Baseline

The second baseline model we trained was a generative adversarial network with a feature loss, which is based on the end-to-end reconstruction model from Shen et al. [43]. We used the generator and discriminator modules present in brain2pix, however, we did not construct RFSimages for the input of the model. Instead, this baseline model takes in raw fMRI data as input signals, and thus does not exploit the topographical information of the visual cortex and the spatial nature of the stimulus.

Reconstructions of this method is shown in Figure 3 below “Baselines > Shen et al. > Rec.”. Although we trained both the models with the same amount of epochs and the same amount of data, quantitative and qualitative results show that our model outperforms the Shen et. al. baseline model (see Table 1 and Table 2. For instance, some static black and white noise is present in the reconstructions of row 2, 4, 6. Additionally, the Ood character is not recognizable. This model appeared to perform better than the Nishimoto et al. baseline on the Doctor Who dataset.

Our analyses indicates that our method outperforms both baselines. Implementation details can be found in the source codes of these baselines which are included in the supplementary material in the github repository.

### ROI experiments

In order to isolate the role of the regions of interest, we performed a series of follow up experiments where only data from one ROI was provided to the network. We used a fixed receptive field matrix (Eq. 1). All the experimental details are identical to the main experiment. Figure 4 shows the ROI-specific reconstructions. Reconstructions based on V1 tend to have sharper pixelwise correspondence, whereas some high-level features, such as global illumination, were not captured very well. The combined brain2pix model with all ROIs (V1-V3) on the other hand was able to capture the color profile of the scenes very well. These two models generated images that captured further readily interpretable high-level information such as the existence of a person in the scene and even the expressions on the faces of individuals in the scenes. It is interesting to note that the ROIs did not contain higher-level brain regions, such as lateral occipital cortex that play a large role in object perception and fusiform face area that specializes in face processing.

**Figure 4.**
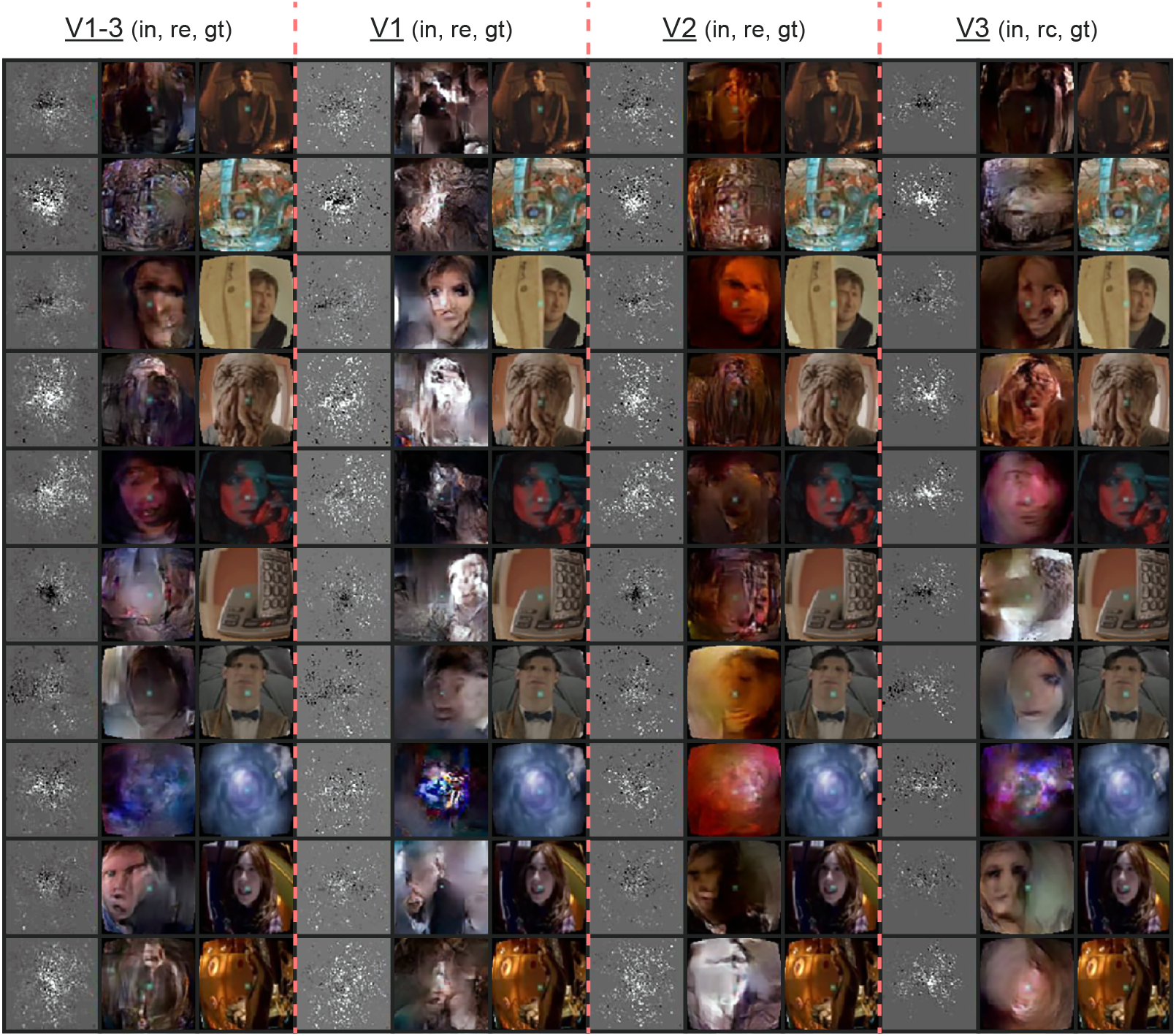
ROI experiment: Reconstructions from the brain2pix FixedRF method trained on various brain regions. Columns 1–3 show the inputs (in), reconstructions (re) and ground truths (gt) of all combined regions (V1–V3), respectively. Columns 4–6 show these results for only V1, columns 7–9 for V2, and finally columns 10–12 are results belonging to the model trained only on V3.

The quantitative results are given in Table 3 and Table 4. The combined model performs substantially better than the individual models with the V1 model having the worst performance. V2 and V3 were similar to each other in quantitative performance. The poor performance of V1 is likely due to the fact that the adversarial and feature losses had a larger weight in the training process, biasing the model towards using higher-order features for reconstruction.

**Table 3.** ROI experiment *correlation values*: Correlation between Alexnet and C3D features for the ROI experiment. V1, V2, V3 are the individual regions of interests and V1–V3 are all three visual layers combined.

|  | V1-3 | V1 | V2 | V3 |
| --- | --- | --- | --- | --- |
| AlexNet pool2 | **0.4615** | 0.2626 | 0.4247 | 0.4178 |
| AlexNet pool5 | **0.3507** | 0.1920 | 0.3387 | 0.3294 |
| AlexNet fc6 | **0.4608** | 0.2704 | 0.4453 | 0.4304 |
| C3D pool2 | **0.4868** | 0.2640 | 0.4704 | 0.4685 |
| C3D pool5 | **0.2426** | 0.1299 | 0.2224 | 0.2157 |
| C3D fc6 | **0.2519** | 0.1349 | 0.2302 | 0.2228 |

**Table 4.** ROI experiment *distance values*: Distances between Alexnet and C3D features for the ROI experiment. V1, V2, V3 are the individual regions of interests (ROIs) and V1–V3 are all three ROIs combined.

|  | V1-3 | V1 | V2 | V3 |
| --- | --- | --- | --- | --- |
| AlexNet pool2 | **4.6252** | 5.9592 | 4.8401 | 5.1095 |
| AlexNet pool5 | 1.2691 | 1.6985 | **1.2673** | 1.2898 |
| AlexNet fc6 | **0.0899** | 2.0553 | 1.4820 | 1.4877 |
| C3D pool2 | 1.3983 | 1.8996 | **1.3949** | 1.4288 |
| C3D pool5 | **3.4626** | 4.5135 | 3.5756 | 3.5542 |
| C3D fc6 | **0.9332** | 1.1592 | 0.9873 | 0.9417 |

### Ablation experiments

The ablation studies were performed to test the impact of VGG-loss and adversarial loss on the performance of the model (see Table 5 and Table 6). “No adversarial” refers to the brain2pix without a discriminator loss, using only the VGG-feature loss to optimize the model. In this ablation case, the model did not learn to reconstruct images but rather outputted square patterns that repeated across all images. The second model is the “no feature” model, which was trained without the VGG-loss, thus only making use of the adversarial loss. This resulted in images that look like reconstructions but did not approximate the target.

**Table 5.** Ablation experiment *correlations values.* Correlation between Alexnet and C3D features of the test stimuli and their reconstructions. The brain2pix is compared with two models with either no feature loss or no adversarial loss.

|  | brain2pix | No feature | No adversarial |
| --- | --- | --- | --- |
| AlexNet pool2 | **0.4615** | 0.3950 | 0.1626 |
| AlexNet pool5 | **0.3507** | 0.3168 | 0.1301 |
| AlexNet fc6 | **0.4608** | 0.4273 | 0.1567 |
| C3D pool2 | **0.4868** | 0.4396 | 0.16286 |
| C3D pool5 | **0.2426** | 0.2123 | 0.0396 |
| C3D fc6 | **0.2519** | 0.2255 | 0.0495 |

**Table 6.** Ablation experiment *distance values.* Distances between reconstruction and target features of various layers obtained from passing through a pretrained network (Alexnet and C3D). Here comparisons are done for the brain2pix models and ablation models.

|  | brain2pix | No feature | No adversarial |
| --- | --- | --- | --- |
| AlexNet pool2 | **4.6252** | 5.6450 | 13.1021 |
| AlexNet pool5 | **1.2691** | 1.4201 | 1.5935 |
| AlexNet fc6 | **0.0899** | 1.5991 | 2.1505 |
| C3D pool2 | **1.3983** | 1.5988 | 7.5023 |
| C3D pool5 | **3.4626** | 3.6445 | 5.6622 |
| C3D fc6 | **0.9332** | 0.9657 | 2.7328 |

### Experiment with various data size

Finally, we wanted to see how the model performs when less data is fed into the network. This was done by training our model on a selected amount of Doctor Who frames from the training set. The full training set, which was worth ≈ 24 hours of data was split into the following: 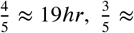 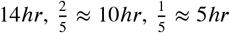 which resulted in training sizes of 95.200, 71.400, 37.600, and 23.800 frames with 50 epochs for each condition. The reconstructions of the splitted data is then compared with the full ~ 24hr worth of data (also trained on 50 epochs). The reconstructions of this experiment are shown in Figure 5. Although the highest amount of data shows the best reconstructions, we still see notable reconstructions based on smaller amounts of data, which suggests that it is not necessary to record 23 hours worth of data to achieve reconstructions with the brain2pix model. The quantitative results can be found in Table 7 and Table 8.

**Figure 5.**
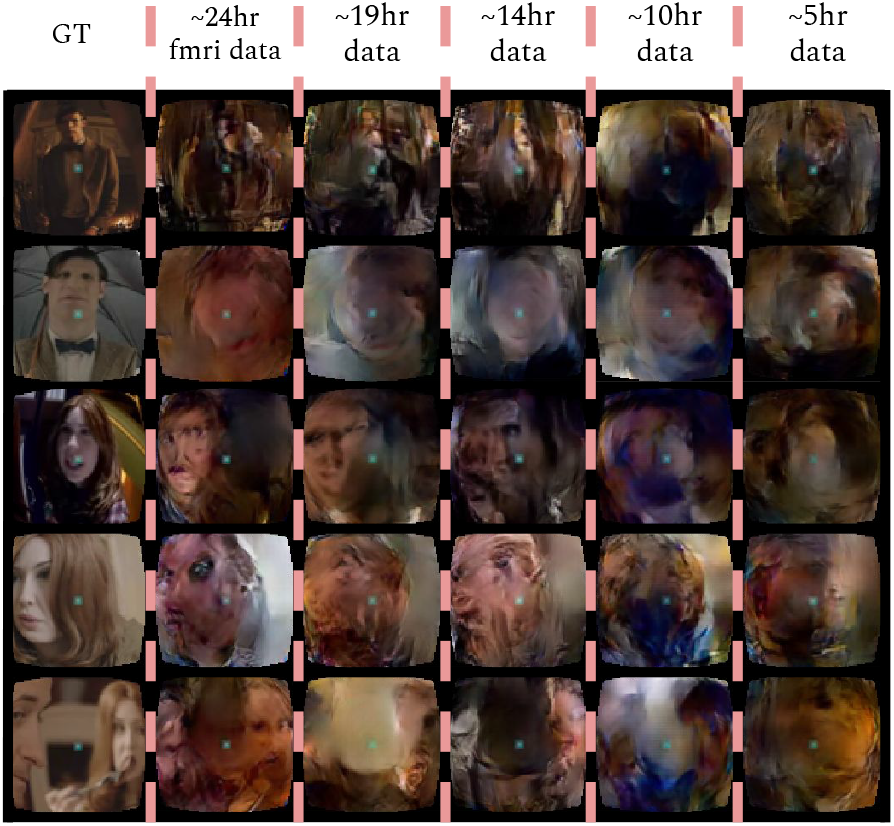
Time experiment: Reconstructions from the brain2pix fixed RF method trained on various amounts of fMRI data after 50 epochs of training. Ground truth (GT), ~24hr is the entire dataset, 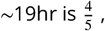, 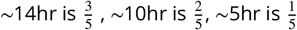 of the dataset.

**Table 7.** Splitted data *correlation* values: the model was trained on data worth of 5, 10, 14, 19, and 24 hours.

| | B2P ($\sim 24$h) | B2P ($\sim 19$h) | B2P ($\sim 14$h) | B2P ($\sim 10$h) | B2P ($\sim 5$h) |
| --- | --- | --- | --- | --- | --- |
| AlexNet pool2 | **0.4615** | 0.4388 | 0.4262 | 0.4132 | 0.4342 |
| AlexNet pool5 | **0.3507** | 0.3289 | 0.3236 | 0.3195 | 0.3206 |
| AlexNet fc6 | **0.4608** | 0.4399 | 0.4270 | 0.4339 | 0.4376 |
| C3D pool2 | **0.4869** | 0.4797 | 0.4270 | 0.4558 | 0.4714 |
| C3D pool5 | **0.2426** | 0.2037 | 0.1933 | 0.2061 | 0.2057 |
| C3D fc6 | **0.2519** | 0.2171 | 0.2061 | 0.2325 | 0.2226 |

**Table 8.**
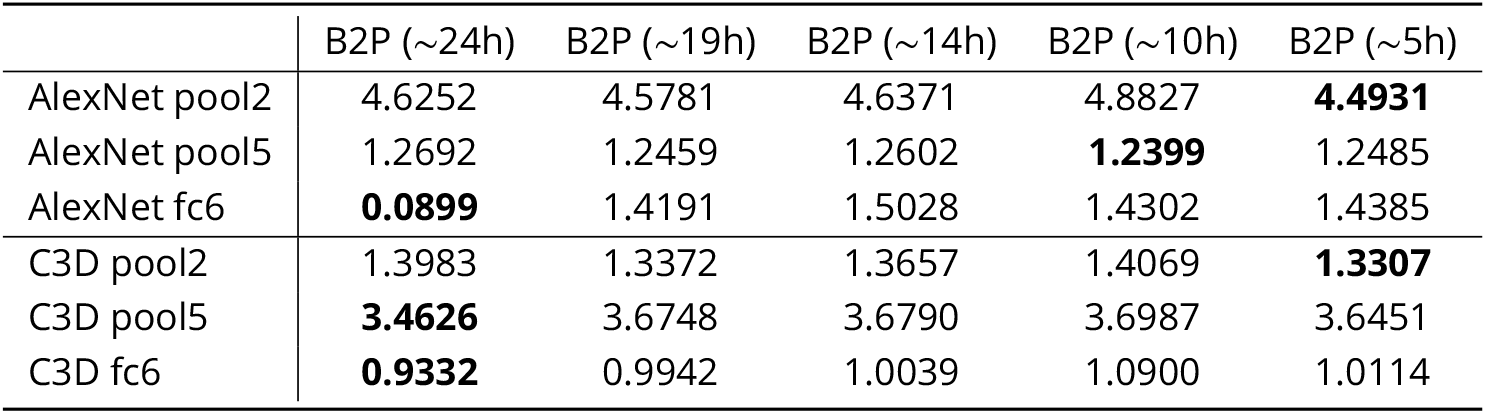
Splitted data *distance* values: the model was trained on data worth of 5, 10, 14, 19, and 24 hours.

## Discussion

In this paper, we introduced a new brain reconstruction method, brain2pix, which exploits the topographic organization of the visual cortex by mapping brain activation to a linear pixel space where it is then processed with a fully convolutional image-to-image network. To the best of our knowledge, this is the first end-to-end approach capable of generating semantically accurate reconstructions from a naturalistic continuous video stream. Furthermore, our approach was shown to outperform other baseline decoding methods.

One of the next challenges that we can try to tackle is decoding at higher frame rates rather than using the same number of frames as brain signals in order to reconstruct even finer temporal details. Additionally, in our current experiments we focused mainly on optimizing our model based on responses from the early visual regions V1, V2, V3. A natural extension of the current work is to extend our focus to the higher level areas in the temporal and parietal cortex. Since these areas process coarse-grained semantic information, experiments feeding their responses to deeper layers of the network could reveal reconstructions driven by semantics. We can also apply the model on imagery data.

Neural decoding studies are crucial for understanding the functioning of the human brain, broadly benefiting the field of neuroscience. Furthermore, neural decoding algorithms make up a major component of brain-computer interfaces (BCIs). Brain-computer interfaces enable disabled people to perform tasks that they would not be able to perform otherwise, by substituting their lost faculties. These technologies can range from a communication interface for a locked-in patient, to a neuroprosthetic limb, and more. While the algorithms that we develop and study in this paper are specialized to reconstruct visual stimuli from brain responses, we foresee that the suggested principles can be applied to different applications, with some adaptations. For instance, we use a relatively slow signal (BOLD response), which reflects the neural responses that take place several seconds prior to them. A time-critical BCI system would need to make use of a signal with no such delays to perform well.

While admittedly the promise of algorithms that reconstruct internally generated or externally induced percepts is yet to be fully achieved, scientists that attempt to extract information from the brain should ensure the safety and privacy of the users [19]. Future studies should make sure to follow similar strict regulations, ensuring only a positive impact of these fascinating methods that allow us to peek into the human mind.

## Notes

https://github.com/Neural-Coding/Brain2Pix

### Competing Interest Statement

The authors have declared no competing interest.

## References

[1] Bäckström K, Nazari M, Gu IY, Jakola AS. An efficient 3D deep convolutional network for Alzheimer’s disease diagnosis using MR images. In: 2018 IEEE 15th International Symposium on Biomedical Imaging IEEE; 2018. p. 149–153.

[2] Bashivan P, Kar K, DiCarlo JJ. Neural population control via deep image synthesis. Science. 2019; 364(6439):9436.

[3] Chen T, Li M, Li Y, Lin M, Wang N, Wang M, Xiao T, Xu B, Zhang C, Zhang Z. MXNet: A Flexible and Efficient Machine Learning Library for Heterogeneous Distributed Systems.. 2015.

[4] Cohen TS, Geiger M, Köhler J, Welling M. Spherical CNNs. In: International Conference on Learning Representations; 2018.

[5] Dado T, Gucluturk Y, Ambrogioni L, Ras G, Bosch SE, van Gerven M, Guclu U. Hyperrealistic neural decoding: Linear reconstruction of face stimuli from fMRI measurements via the GAN latent space. bioRxiv. 2020.

[6] Dong C, Loy CC, Tang X. Accelerating the super-resolution convolutional neural network. In: European Conference on Computer Vision Springer; 2016. p. 391–407.

[7] Dumoulin SO, Wandell BA. Population receptive field estimates in human visual cortex. Neuroimage. 2008; 39(2):647–660.

[8] Fey M, Eric L J, Weichert F, Müller H. SplineCNN: Fast geometric deep learning with continuous B-spline kernels. In: Proceedings of the IEEE Conference on Computer Vision and Pattern Recognition; 2018. p. 869–877.

[9] Fischl B. FreeSurfer. Neuroimage. 2012; 62(2):774–781.

[10] van Gerven MA, de Lange FP, Heskes T. Neural decoding with hierarchical generative models. Neural Computation. 2010; 22(12):3127–3142.

[11] Güçlütürk Y, Güçlü U, van Lier R, van Gerven MA. Convolutional sketch inversion. In: European Conference on Computer Vision Springer; 2016. p. 810–824.

[12] Güçlütürk Y, Güçlü U, Seeliger K, Bosch S, van Lier R, van Gerven MA. Reconstructing perceived faces from brain activations with deep adversarial neural decoding. In: Advances in Neural Information Processing Systems; 2017. p. 4246–4257.

[13] Han K, Wen H, Shi J, Lu KH, Zhang Y, Fu D, Liu Z. Variational autoencoder: An unsupervised model for encoding and decoding fMRI activity in visual cortex. NeuroImage. 2019 sep; 198:125–136. https://doi.org/10.1016%2Fj.neuroimage.2019.05.039, doi: 10.1016/j.neuroimage.2019.05.039.

[14] Henschen SE. On the visual path and centre. Brain. 1893; 16(1-2):170–180.

[15] Holmes G, Lister W. Disturbances of vision from cerebral lesions, with special reference to the cortical representation of the macula. Brain. 1916; 39(1-2):34–73.

[16] Horikawa T, Kamitani Y. Hierarchical Neural Representation of Dreamed Objects Revealed by Brain Decoding with Deep Neural Network Features. Frontiers in Computational Neuroscience. 2017; 11:4. https://www.frontiersin.org/article/10.3389/fncom.2017.00004, doi: 10.3389/fncom.2017.00004.

[17] Horikawa T, Tamaki M, Miyawaki Y, Kamitani Y. Neural decoding of visual imagery during sleep. Science. 2013; 340(6132):639–642.

[18] Hubel DH, Wiesel TN. Receptive fields, binocular interaction and functional architecture in the cat’s visual cortex. The Journal of Physiology. 1962; 160(1):106–154.

[19] Ienca M, Haselager P, Emanuel EJ. Brain leaks and consumer neurotechnology. Nature Biotechnology. 2018; 36(9):805–810. doi: 10.1038/nbt.4240.

[20] Iizuka S, Simo-Serra E, Ishikawa H. Let there be color! Joint end-to-end learning of global and local image priors for automatic image colorization with simultaneous classification. ACM Transactions on Graphics. 2016; 35(4):1–11.

[21] Inouye T. Die Sehstorungen bei Schussverletzungen der kortikalen Sehsphare. Nach Beobachtungen an Verwundeten der letszten Japanischen Kriege. 1909.

[22] Isola P, Zhu JY, Zhou T, Efros AA. Image-to-image translation with conditional adversarial networks. In: Proceedings of the IEEE Conference on Computer Vision and Pattern Recognition; 2017. p. 1125–1134.

[23] Kay W, Carreira J, Simonyan K, Zhang B, Hillier C, Vijayanarasimhan S, Viola F, Green T, Back T, Natsev P, et al. The kinetics human action video dataset. arXiv preprint arXiv:170506950. 2017;.

[24] Kim J, Kwon L J, Mu L K. Deeply-recursive convolutional network for image super-resolution. In: Proceedings of the IEEE Conference on Computer Vision and Pattern Recognition; 2016. p. 1637–1645.

[25] Kok P, Jehee JF, De Lange FP. Less is more: expectation sharpens representations in the primary visual cortex. Neuron. 2012; 75(2):265–270.

[26] Kondor R, Lin Z, Trivedi S. Clebsch–gordan nets: a fully Fourier space spherical convolutional neural network. In: Advances in Neural Information Processing Systems; 2018. p. 10117–10126.

[27] Krizhevsky A, Sutskever I, Hinton GE. Imagenet classification with deep convolutional neural networks. In: Advances in Neural Information Processing Systems; 2012. p. 1097–1105.

[28] Li Y, Qi H, Dai J, Ji X, Wei Y. Fully convolutional instance-aware semantic segmentation. In: Proceedings of the IEEE Conference on Computer Vision and Pattern Recognition; 2017. p. 2359–2367.

[29] Long J, Shelhamer E, Darrell T. Fully convolutional networks for semantic segmentation. In: Proceedings of the IEEE Conference on Computer Vision and Pattern Recognition; 2015. p. 3431–3440.

[30] Miyawaki Y, Uchida H, Yamashita O, Sato Ma, Morito Y, Tanabe HC, Sadato N, Kamitani Y. Visual image reconstruction from human brain activity using a combination of multiscale local image decoders. Neuron. 2008; 60(5):915–929.

[31] Monti F, Boscaini D, Masci J, Rodola E, Svoboda J, Bronstein MM. Geometric deep learning on graphs and manifolds using mixture model CNNs. In: Proceedings of the IEEE Conference on Computer Vision and Pattern Recognition; 2017. p. 5115–5124.

[32] Naselaris T, Prenger RJ, Kay KN, Oliver M, Gallant JL. Bayesian reconstruction of natural images from human brain activity. Neuron. 2009; 63(6):902–915.

[33] Nishimoto S, Gallant JL. A three-dimensional spatiotemporal receptive field model explains responses of area MT neurons to naturalistic movies. Journal of Neuroscience. 2011; 31(41):14551–14564. doi: 10.1523/JNEUROSCI.6801-10.2011.

[34] Nishimoto S, Vu AT, Naselaris T, Benjamini Y, Yu B, Gallant JL. Reconstructing visual experiences from brain activity evoked by natural movies. Current Biology. 2011; 21(19):1641–1646.

[35] Noh H, Hong S, Han B. Learning deconvolution network for semantic segmentation. In: Proceedings of the IEEE International Conference on Computer Vision; 2015. p. 1520–1528.

[36] Ronneberger O, Fischer P, Brox T. U-net: Convolutional networks for biomedical image segmentation. In: International Conference on Medical Image Computing and Computer-Assisted Intervention Springer; 2015. p. 234–241.

[37] Russakovsky O, Deng J, Su H, Krause J, Satheesh S, Ma S, Huang Z, Karpathy A, Khosla A, Bernstein M, Berg AC, Fei-Fei L. ImageNet Large Scale Visual Recognition Challenge. International Journal of Computer Vision (IJCV). 2015; 115(3):211–252. doi: 10.1007/s11263-015-0816-y.

[38] Sarraf S, Tofighi G. Classification of Alzheimer’s disease using fMRI data and deep learning convolutional neural networks. arXiv preprint arXiv:160308631. 2016;.

[39] Seeliger K, Ambrogioni L, Güçlü U, van Gerven M. End-to-end neural system identification with neural information flow. PLoS Computational Biology. 2021 02; 17:1–22.

[40] Seeliger K, Sommers R, Güçlü U, Bosch S, van Gerven M. A large single-participant fMRI dataset for probing brain responses to naturalistic stimuli in space and time. BioRxiv. 2019; p. 687681.

[41] Seeliger K, Güçlü U, Ambrogioni L, Güçlütürk Y, van Gerven MA. Generative adversarial networks for reconstructing natural images from brain activity. NeuroImage. 2018; 181:775–785.

[42] Selim A, Elgharib M, Doyle L. Painting style transfer for head portraits using convolutional neural networks. ACM Transactions on Graphics. 2016; 35(4):1–18.

[43] Shen G, Dwivedi K, Majima K, Horikawa T, Kamitani Y. End-to-End Deep Image Reconstruction From Human Brain Activity. Frontiers in Computational Neuroscience. 2019; 13:21. https://www.frontiersin.org/article/10.3389/fncom.2019.00021, doi: 10.3389/fncom.2019.00021.

[44] Shen G, Horikawa T, Majima K, Kamitani Y. Deep image reconstruction from human brain activity. PLoS Computational Biology. 2019 jan; 15(1):e1006633. https://doi.org/10.1371%2Fjournal.pcbi.1006633, doi: 10.1371/journal.pcbi.1006633.

[45] Thirion B, Duchesnay E, Hubbard E, Dubois J, Poline JB, Lebihan D, Dehaene S. Inverse retinotopy: inferring the visual content of images from brain activation patterns. Neuroimage. 2006; 33(4):1104–1116.

[46] Tran D, Bourdev L, Fergus R, Torresani L, Paluri M. Learning spatiotemporal features with 3D convolutional networks. In: Proceedings of the IEEE International Conference on Computer Vision; 2015. doi: 10.1109/ICCV.2015.510.

[47] Wen H, Shi J, Zhang Y, Lu KH, Cao J, Liu Z. Neural Encoding and Decoding with Deep Learning for Dynamic Natural Vision. Cerebral Cortex. 2017 oct; 28(12):4136–4160. https://doi.org/10.1093%2Fcercor%2Fbhx268, doi: 10.1093/cercor/bhx268.

[48] Zhang R, Isola P, Efros AA. Colorful image colorization. In: European Conference on Computer Vision Springer; 2016. p. 649–666.

[49] Zhang R, Zhu J, Isola P, Geng X, Lin AS, Yu T, Efros AA. Real-time user-guided image colorization with learned deep priors. arXiv preprint arXiv:170502999. 2017.

[50] Zhang Y, Qiu Z, Yao T, Liu D, Mei T. Fully convolutional adaptation networks for semantic segmentation. In: Proceedings of the IEEE Conference on Computer Vision and Pattern Recognition; 2018. p. 6810–6818.

[51] Zhang Y, Tian Y, Kong Y, Zhong B, Fu Y. Residual dense network for image super-resolution. In: Proceedings of the IEEE Conference on Computer Vision and Pattern Recognition; 2018. p. 2472–2481.

[52] Zhu JY, Park T, Isola P, Efros AA. Unpaired image-to-image translation using cycle-consistent adversarial networks. In: Proceedings of the IEEE International Conference on Computer Vision; 2017. p. 2223–2232.

